# Solid-state NMR of paired helical filaments formed by the core tau fragment tau(297-391)

**DOI:** 10.1101/2022.06.09.495520

**Authors:** Youssra K. Al-Hilaly, Connor Hurt, Janet E. Rickard, Charles R. Harrington, John M.D. Storey, Claude M. Wischik, Louise C. Serpell, Ansgar B. Siemer

## Abstract

Aggregation of the tau protein into fibrillar cross-β aggregates is a hallmark of Alzheimer’s diseases (AD) and many other neurodegenerative tauopathies. Recently, several core structures of patient-derived tau paired helical filaments (PHFs) have been solved revealing a structural variability that often correlates with a specific tauopathy. To further characterize the dynamics of these fibril cores, to screen for strain-specific small molecules as potential biomarkers and therapeutics, and to develop strain-specific antibodies, recombinant in-vitro models of tau filaments are needed. We recently showed that a 95-residue fragment of tau (from residue 297 to 391), termed dGAE, forms filaments in vitro in the absence of polyanionic co-factors often used for in vitro aggregation of full length tau. Tau(297-391) was identified as the proteolytic resistant core of tau PHFs and overlaps with the structures characterized by cryo-electron microscopy in ex-vivo PHFs, making it a promising model for the study of AD tau filaments in vitro. In the present study, we used solid-state NMR to characterize tau(297-391) filaments and show that such filaments assembled under non-reducing conditions are more dynamic and less ordered than those made in the presence of the reducing agent, DTT. We further report the resonance assignment of tau(297-392)+DTT filaments and compare it to existing core structures of tau.

## Introduction

Tau accumulates intracellularly in neurofibrillary tangles as paired helical and straight filaments in AD and tau also accumulates as filaments in several other neurodegenerative diseases collectively known as tauopathies (Vaquer-Alicea et al., 2021). The presence of tau filaments is strongly correlated with dementia in tauopathies (Wilcock and Esiri, 1982; Arriagada et al., 1992; Mukaetova-Ladinska et al., 2000; Maruyama et al., 2013; Brier et al., 2016; Gomperts et al., 2016; Smith et al., 2016) and oligomeric tau species are implicated in the spread of tau aggregates through the brain (Clavaguera et al., 2009; Sanders et al., 2014). Tau is encoded by the *MAPT* gene, which produces six isoforms in the human nervous system through alternative splicing (Goedert et al., 1989). These isoforms range from 352 to 441 amino acid residues and contain either three (3R) or four (4R) tandem imperfect repeats, which are 31 of 32 residues in length (Crowther et al., 1989). Monomeric tau is able to stabilize microtubules and is also thought to play roles in signal transduction, actin interaction, and the binding of pericentromeric chromatin in the nucleus (Buée et al., 2000; Bukar Maina et al., 2016; Mansuroglu et al., 2016).

Early studies on tau filaments showed that they form canonical cross-β (amyloid) structures with an in-register parallel-β core (Berriman et al., 2003; Margittai and Langen, 2004). In recent years, structures of the cross-β cores of tau filaments extracted from patient tissue have been resolved thanks to advances in cryo-electron microscopy (cryo-EM). These structures showed that specific tauopathies such as corticobasal degeneration (CBD) have unique tau fibril cores, whereas filaments extracted from other diseases showed fibril core structures very similar to those found in AD (Fitzpatrick et al., 2017; Hallinan et al., 2021; Shi et al., 2021). This phenomenon of disease-specific fibril strains has been observed for other cross-β fibrils in disease including those formed by α-synuclein (Bousset et al., 2013; Mehra et al., 2021; Sawaya et al., 2021).

Despite advances in our understanding of patient-derived tau filaments, there is a need for in vitro models of these filaments to facilitate the screening of small molecules as potential biomarkers or therapeutics and allow the production of strain-specific antibodies.

Full-length tau is highly soluble and not very aggregation prone. Therefore, heparin or other polyanionic molecules have been used to induce aggregation in vitro (Goedert et al., 1996; Kampers et al., 1999; Pérez et al., 2001). However, a truncated tau (297-391), first recognised as the protease-resistant core of PHFs in AD (Jakes et al., 1991; Novak et al., 1993) was found to form filaments in vitro in the absence of any additives (Al-Hilaly et al., 2017). This fragment of tau is also termed dGAE, where ‘d’ corresponds to the amino acid Ile-297 and ‘GAE’ to the three C-terminal amino acids Gly-389, Ala-390, Glu-391 from the 441-residue tau isoform containing 2 N-terminal inserts and 4 tandem repeats (2N4R). Furthermore, tau(297-391) overlaps with the core of filaments resolved from ex-vivo filaments by cryo-EM (Fitzpatrick et al., 2017), was shown to form PHF that resemble those observed in AD patients (Al-Hilaly et al., 2020; Lutter et al., 2022) and later confirmed by cryo-EM to form an architecture identical to those from AD brain under certain conditions (Lövestam et al., 2022). Tau(297-391) has been shown to recruit endogenous tau in a cell model (Harrington et al., 2015) and is able to self-propagate by binding full-length tau reproducing the protease-resistant core following treatment with proteases (Wischik et al., 1996). The tau(297-391) sequence contains a single cysteine residue at position 322 and the oxidation/reducing conditions of assembly have been shown to have a profound effect on the assembly and morphology of the resulting filaments due to the formation of a disulphide bound dimer in under non-reducing conditions (Al-Hilaly et al., 2017).

Here we examine the consequences of oxidation conditions on the structure and dynamics of the tau(297-391) filaments and compare the structure of such filaments to ex-vivo filaments from AD patients resolved by cryo-EM. Our results show that the filaments formed from tau(297-391) are well ordered as long as C322 is reduced. The comparison of our resonance assignment of tau(297-391) filaments formed in reducing conditions with cryo-EM structures of tau generally confirms that these filaments share the cross-β core structure with AD type ex vivo filaments.

## Materials and Methods

### Protein expression, purification, and filament formation

Uniformly ^13^C-^15^N labeled tau(297-391) was expressed in *E. coli* following a protocol described by Marley and co-workers (Marley et al., 2001). The purification was done as previously described (Al-Hilaly et al., 2017, 2020). To initiate filament formation, 300 μM of ^13^C-^15^N labeled tau(297-391) was incubated in phosphate buffer (10 mM, pH 7.4) with and without 1,4 dithiothreitol (DTT) (10 mM) and agitated at 400 oscillations per minute (Eppendorf Thermomixer C, Eppendorf, Germany) for 48 h.

### NMR experiments and data analysis

Uniformly ^13^C-^15^N-labeled tau(297-391) filaments were centrifuged into 1.6 mm rotors using a home built packing tool similar to the one described by Böckmann and co-workers (Böckmann et al., 2009). The following experimental details apply to all NMR experiments: an Agilent DD2 spectrometer with a proton frequency of 600 MHz equipped with a 1.6 mm triple-resonance MAS probe was used. The set-temperature for all experiments was 0°C. ^1^H, ^13^C, ^15^N hard pulses were done with rf-field strengths of 200, 100, and 50 kHz, respectively. High-power ^1^H decoupling was done using the XiX decoupling scheme (Detken et al., 2002) during direct and indirect time domains and CW decoupling was applied during DREAM and double CP recoupling steps all with a ^1^H rf-field strength of 140 kHz. 1D cross polarization (CP) and refocused INEPT experiments were recorded at 25 kHz MAS, a spectral width of 50 kHz and 1024 acquisitions were added for each of these spectra. The initial ^1^H-^13^C CP was done with a contact time of 1 ms and rf-field strengths of 85 and 60 kHz on ^1^H and ^13^C, respectively. The 2D PARIS experiment used the same CP to create the initial ^13^C magnetization followed by a 500 ms of PARIS ^13^C mixing using a 10 kHz recoupling field on ^1^H and the N=0.5 condition (Weingarth et al., 2009). Spectral widths were 50 kHz in both dimensions and 112 acquisitions were co-added for each of the 350 complex, indirect increments. The 2D DREAM was recorded at 35 kHz MAS with 4.5 ms of DREAM recoupling and 32 acquisitions were co-added for each of the 800 real, TPPI increments. The 2D NCA spectrum was recorded at 35 kHz MAS using a standard double CP (DCP) pulse sequence using an initial 1.5 ms ^1^H-^15^N CP with rf-field strengths of 85 kHz on ^1^H and 50 kHz on ^15^N followed by 8 ms of SPECIFIC CP (Baldus et al., 1998) with rf-field strengths of 40 kHz and 5 kHz on ^15^N and ^13^C, respectively, and the ^13^C transmitter set to 52 ppm. Spectral widths were 50 kHz for ^13^C and 3 kHz for ^15^N and 128 acquisitions were co-added for each of the 40 complex, indirect increments. The NCO spectrum was acquired with the same rf-field strengths but the ^13^C transmitter set to 174 ppm instead and a mixing time of 9 ms. In addition, homonuclear decoupling using the LOW BASHD scheme (Struppe et al., 2013) was applied during t_1_. Spectral widths were 2.5 kHz for ^13^C and 3 kHz for ^15^N and 96 acquisitions were co-added for each of the 40 complex, indirect increments.

The NCOCA was recorded at 35 kHz MAS using a double CP followed by a DREAM sequence for homonuclear ^13^C-^13^C transfer (Detken et al., 2001). The ^1^H-^15^N and ^15^N-^13^C CPs were the same as for the NCO 2D followed by 2.5 ms of DREAM recoupling at the HORROR condition of 17.5 kHz during which the ^13^C transmitter was moved to 115 ppm. Spectral widths were 50 kHz for the direct and 3 kHz both of the indirect dimensions, each of which was sampled with 32 indirect, complex points and 64 acquisitions per increments that were recorded using non-uniformly sampling with 25% coverage. The NCACO was recorded with the same pulse sequence as the NCOCA spectrum but using the double CP conditions used for the 2D NCA spectrum and the same DREAM recoupling conditions. Spectral widths were 50 kHz and 5 kHz for the direct and indirect ^13^C dimensions, and 3 kHz for the ^15^N dimension. Thirty two acquisitions were co-added for each of the 32 indirect ^13^C and 20 indirect ^15^N increments that were acquired using non-uniformly sampling at 25% coverage.

The NCACB 3D was recorded at 25 kHz MAS using the same pulse sequence as the NCOCA spectrum. ^1^H and ^15^N rf-field strengths of 65 kHz and 40 kHz were used for the 1st CP with a contact time of 1.5 ms and ^15^N and ^13^C rf-field strengths of 32.5 kHz and 7.5 kHz for the 2nd CP with a contact time of 9 ms and the ^13^C transmitter set to 45 ppm. DREAM recoupling at the HORROR condition of 12.5 kHz was done for 3.5 ms for the final transfer step. Spectral widths were 50 kHz and 6 kHz for the direct and indirect ^13^C dimensions, and 3 kHz for the ^15^N dimension and 56 acquisitions were co-added for each of the 50 indirect ^13^C and 26 indirect ^15^N increments that were acquired using non-uniformly sampling at 25% coverage.

The 3D CANcoCA experiment (see Fig. S1) was recorded at 35 kHz MAS using three consecutive CPs and a ^13^C-^13^C DREAM transfer step. The initial ^1^H-^13^C CP was done for 1ms using rf-field strengths of 85 kHz and 50 kHz on ^1^H and ^13^C, respectively. The second and third CP were done for 6 ms with rf-field strength of 40 kHz and 5 kHz on ^15^N and ^13^C, respectively. The CA-N transfer was done with the transmitter at 51 ppm and the NCO transfer with the transmitter at 167 ppm. The final DREAM transfer was done for 2.5 ms at the HORROR condition of 17.5 kHz with the transmitter set to 111 ppm. All NMR time domain datasets were processed using the nmrPipe software package including NUS datasets, which were processed using the IST algorithm implemented in the nmrPipe (Delaglio et al., 1995). Spectra were analyzed using the program CARA (Keller, 2004) and plotted using the nmrglue python package (Helmus and Jaroniec, 2013).

## Results

### Tau(297-391) filaments formed under reducing conditions are more ordered and less dynamic

We previously showed that tau(297-391) formed filaments with different morphologies dependent on the presence of DTT as a reducing agent (Al-Hilaly et al., 2017). To follow up on this finding, we asked how assembling the protein in reducing conditions affects the structure and dynamics of the monomeric constituents of tau(297-391) filaments. To answer this question, we prepared uniformly ^13^C-^15^N labeled tau(297-391) filaments with and without the addition of DTT for solid-state NMR analysis (see Materials and Methods). **Fig. 1** shows 1D ^13^C solid-state NMR spectra of both filaments made in the presence and absence of DTT. We used two different solid-state NMR experiments to highlight regions of different dynamics. The cross polarization (CP) experiment is sensitive to relatively static portions of the sample, which in the case of amyloid fibrils is usually the cross-β core and static framing sequences. The refocused INEPT (RINEPT) experiment, on the other hand, only shows signals coming from regions of considerable dynamics (Heise et al., 2005; Siemer et al., 2006a; Matlahov and van der Wel, 2018; Siemer, 2020). Based on these 1D spectra, the differences between the two fibril types were striking: The filaments formed in the presence of DTT give strong and relatively narrow signals in the CP and only few weak signals in the RINEPT spectrum, indicating that the majority of the tau(297-391) residues are part of a well ordered, static fibril core. In contrast, the filaments formed without DTT give a CP spectrum that is relatively weaker and broader and an RINEPT spectrum that shows many, intense resonances, indicating that these filaments have a less ordered fibril core and that large parts of the sample are in a much more dynamic state.

**Figure 1:**
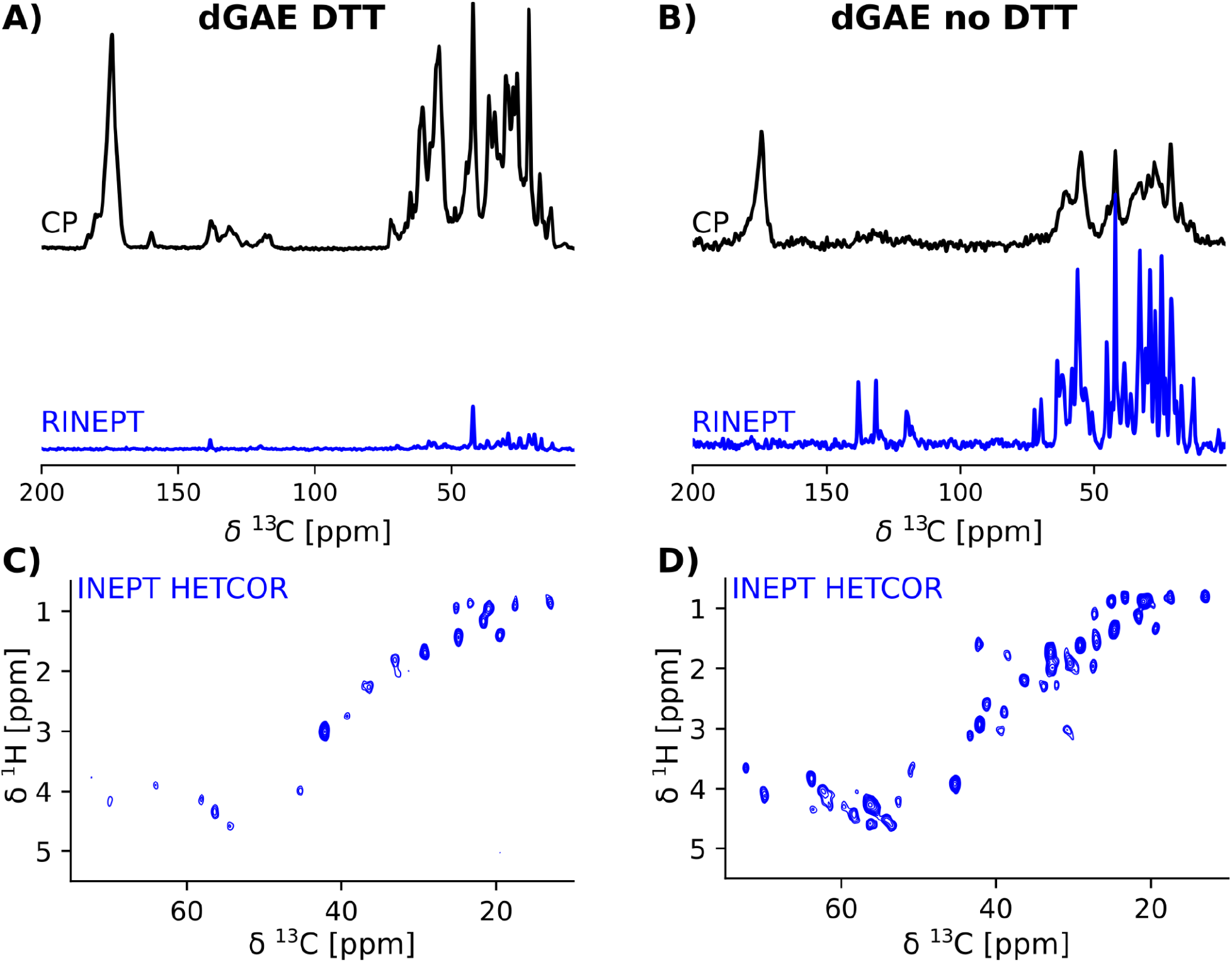
tau(297-391) filaments formed in the presence of DTT are much more rigid and ordered compared to filaments formed without DTT. Solid-state NMR spectra of tau(297-391) filaments formed in the presence (A,C) or absence of DTT (B,D). A,B) 1D ^13^C spectra recorded using ^1^H-^13^C cross polarization (CP, black) and a refocused INEPT transfer (RINEPT, blue) are shown. The CP experiment shows the more static domains of the sample whereas the RINEPT experiment only shows highly dynamic parts of the sample under the conditions used. The strong intensity of the CP spectrum and comparably negligible intensity of the RINEPT spectrum of the filaments made with DTT suggests that the majority of the sample is part of the rigid fibril core. In contrast, the intense RINEPT spectrum of the filaments formed in the absence of DTT suggest that this sample contains a lot of dynamic disorder. C,D) The 2D ^1^H-^13^C INEPT-HETCOR spectra sensitive to dynamic domains confirm this tendency showing many more signals for the sample made in the absence of DTT.

These findings are further supported by the 2D INEPT HETCOR spectra shown on the bottom of **Fig. 1**. These spectra, which are 2D versions of the 1D RINEPT spectrum, show many more and more intense signals for the tau(297-391) filament sample formed without DTT compared to the sample formed in the presence of DTT.

### Resonance assignment of tau(297-391) filaments formed in reducing conditions

Further atomic-resolution NMR analysis of these filaments requires resonance assignments. Because the CP signals of non-DTT tau(297-391) filaments were weak and relatively broad, their assignment was very challenging. Therefore, we focused on tau(297-391) filaments formed in the presence of DTT. To assign the resonances in the static fibril core, we recorded a standard set of ^13^C based solid-state NMR 2D and 3D experiments namely 2D DREAM, NCO, and NCA and 3D NCOCA, NCACB, NCACO, and CANcoCA spectra (Siemer et al., 2006b; Schuetz et al., 2010). We also recorded a ^13^C-^13^C 2D PARIS (Weingarth et al., 2009) spectrum with 500 ms mixing time to confirm some of our assignments.

The 2D DREAM, NCA, and NCO spectra of the tau(297-391)+DTT filaments illustrate the good spectral resolution of this sample (**Fig. 2**) (1D slices through the 2D DREAM are shown in **Fig. S2**). As indicated by the residue-specific labels in **Fig. 2**, we were able to assign the large majority of the cross peaks in these 2D spectra, with the help of the 3D spectra, two of which are illustrated in **Fig. 3**. We identified the following residues in our assignment starting with V309 and Y310 followed by P311-S324, G326-I328, E338-E342, R349-N359, and E372-H374. In addition, we identified a PGGG fragment that is either P332-G335, or P364-G367, or both. The assignment includes backbone ^13^C and ^15^N resonances as well as side chain ^13^C resonances for most of the assigned residues as can be seen from the DREAM spectrum in **Fig. 2A**. The residues of tau(297-391) that were not part of our assignment, could either not be assigned with confidence in our analysis (i.e. the few residues labeled with “?” in **Fig. 2**) or were too dynamic to give intense enough signals in our spectra.

**Figure 2:**
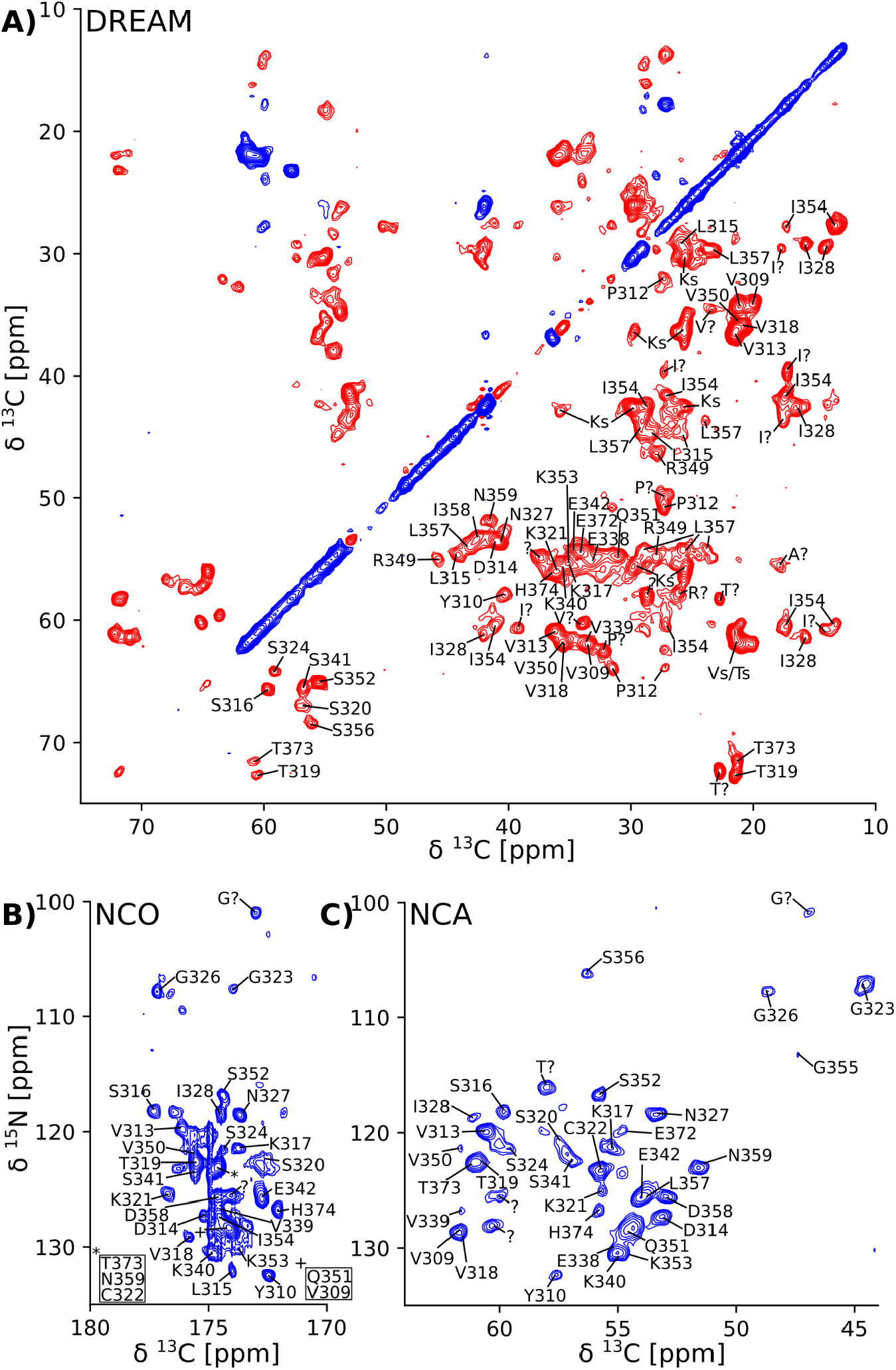
2D solid-state NMR spectra of tau(297-391) filaments made with DTT show narrow lines and one set of cross peaks indicative of a homogenous and well-ordered structure. A) 2D ^13^C-^13^C DREAM spectrum recorded at 35 kHz MAS. Negative and positive cross peaks are shown in red and blue, respectively. B) 2D ^15^N-^13^C NCO spectrum recorded at 35 kHz MAS using LOW BASHD decoupling during t_2_. C) 2D NCA spectrum recorded under conditions similar to the NCO spectrum. Residue-specific resonance assignments are indicated showing an almost complete assignment of these spectra.

**Figure 3:**
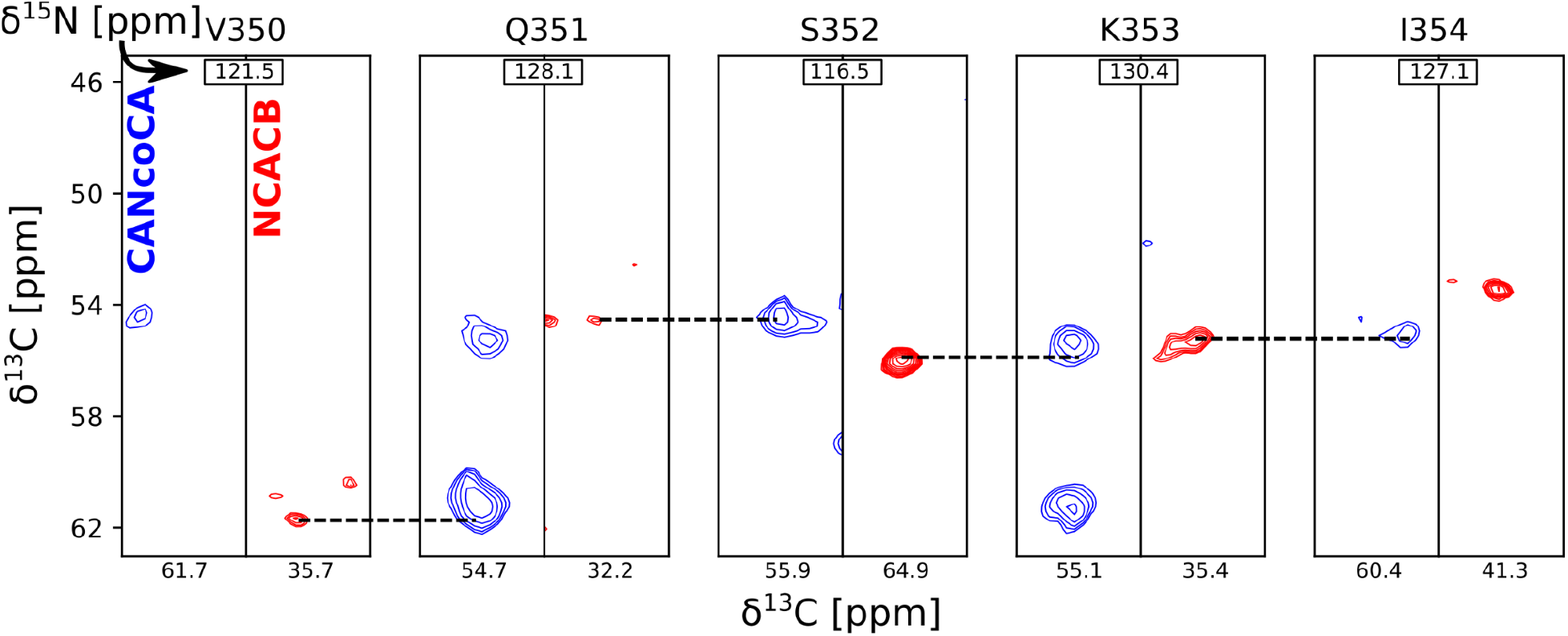
Strip plot of two 3D spectra used for resonance assignment. Strips through 3D CANcoCA (blue) and 3D NCACB (red) spectra recorded at 35 kHz MAS and 25 kHz MAS respectively. The ^15^N shift of each strip is indicated along with the CA (x) and CA-1 (y) shifts for the CANcoCA spectrum and the CB (x) and CA (y) shifts for the NCACB spectrum. Dashed lines indicate the connectivities indicated by the CA-CA-1 peaks.

### Comparison with tau fibril core structures

Because chemical shifts, in particular of Cα and Cβ atoms, are very sensitive to secondary structure (Spera and Bax, 1991), we were now able to ask: How does our solid-state NMR assignment compare to the structure of tau fibril strains isolated from different tauopathies? To address this question, we chose two different approaches: firstly we derived chemical shifts from existing tau fibril core structures and compared them to our measured chemical shifts, and secondly we derived dihedral angles from our NMR assignment and compared them to the dihedral angles of the same tau core structures. For the first approach, we used the program ShiftX2 (Han et al., 2011, 2), which derives chemical shifts from a given PDB file. We focused on Cα and Cβ chemical shifts known to be most sensitive to secondary structure (Spera and Bax, 1991) and examined all tau fibril structures deposited to the PDB, whose cores included the residues found in our solid-state NMR assignment. We then compared the thus calculated chemical shifts (*Cal*) to our NMR assignment (*NMR*) via the sum of the absolute per residue *δCα* and *δCβ* chemical shift difference for every assigned residue *i*

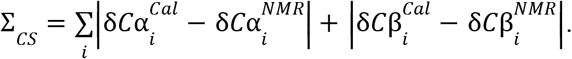

The result is listed in **Table S1** showing that the PDB entry 7mkg i.e. the core of a fibril extracted from a patient with PrP cerebral amyloid angiopathy (Hallinan et al., 2021) resulted in the lowest chemical shift difference. To visualize the per-residue similarity in chemical shift, we calculated the difference between the Cα and Cβ secondary chemical shifts (Δ*S* = Δδ*C*α−Δδ*C*β) for both the chemical shifts derived from the 7mkg structure and our assignment. As can be seen from **Fig. 4A**, there is, with few exceptions, a good agreement in sign and amplitude of the individual Δ*S* values. The sign of Δ*S* is indicative of secondary structure where positive values are found in helical structures and negative values in extended conformations such as β-strands. However, all tau core structures are dominated by β-sheets resulting in negative ΔS values for most residues. What really distinguishes these cores from a secondary structure perspective are the locations of the kinks i.e. the regions that deviate from β-strand conformation. With this in mind, we developed another metric Σ_A_ to compare existing tau core structures with our NMR assignment by calculating the number of residues for which the sign of Δ*S* was different between the two.

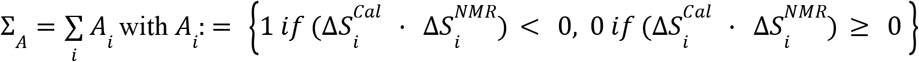

**Figure 4:**
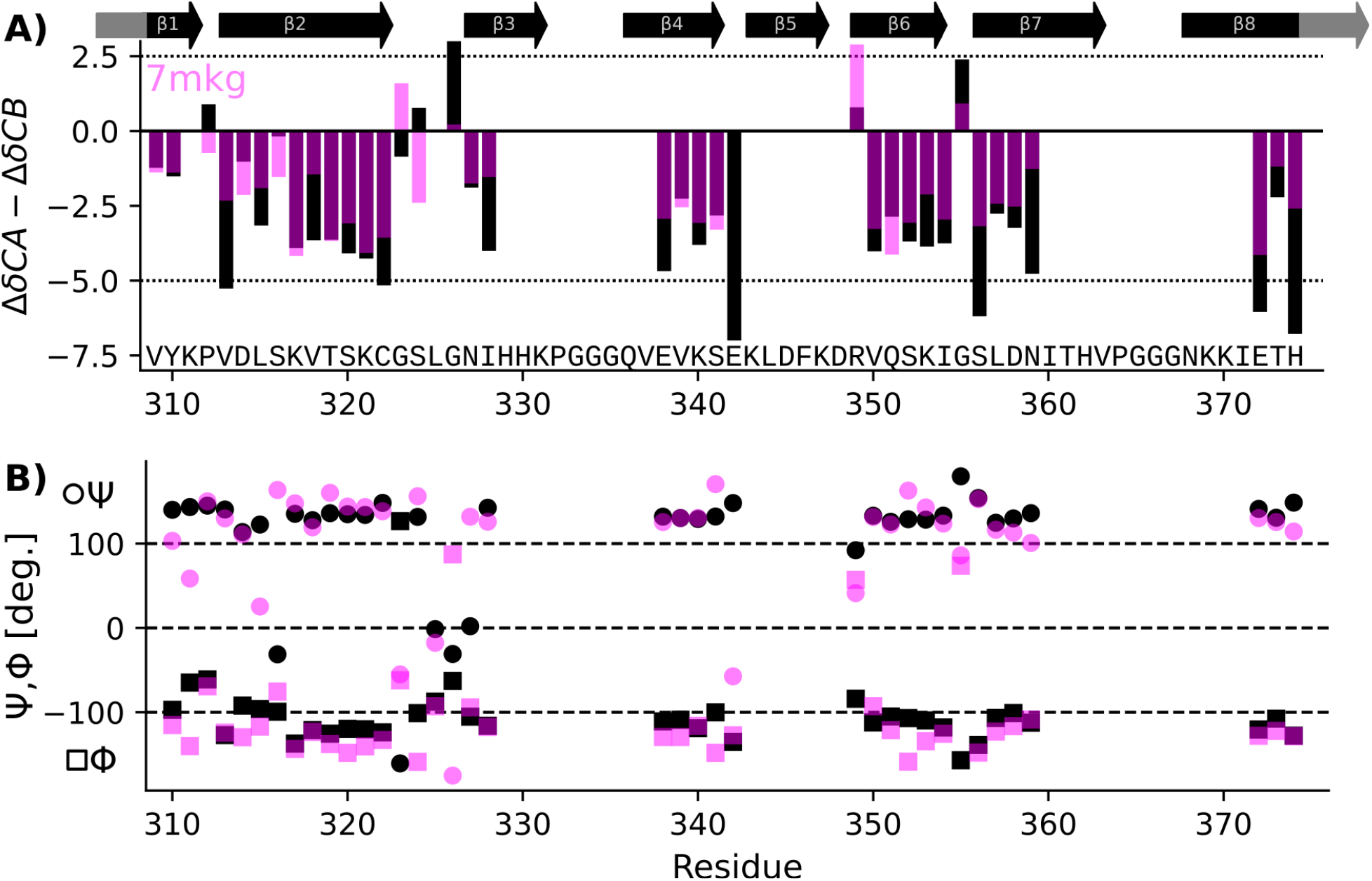
Comparison of NMR assignment and existing cryo-EM structures. A) Difference in CA and CB secondary chemical shift (Δ*S*=ΔδCα-ΔδCβ) as derived from the NMR assignment (black) and calculated from the tau PrP-CAA SF cryo-EM structure (PDB entry 7mkg, fuchsia) using the program ShiftX2. Both secondary shifts overlap well indicating a similar structure. However, there are notable exceptions, specifically P312, G324, S324, G326, and E342. B) Psi (Ψ), phy (Φ) dihedral angles calculated from the NMR assignment using the program

The result of this calculation is shown in **Table S1**. In this case two PDB entries give the best (i.e. lowest) score: 7qjv a structure of recombinant tau(297-391) (Lövestam et al., 2022) and 7p6d the structure of tau fibrils from an argyrophilic grain disease patient (Shi et al., 2021).

For the second approach to compare known tau fibril structures with our NMR assignment, we calculated dihedral Φ,Ψ angles from our assignment (*NMR*) using the program TALOS-N (Shen and Bax, 2013). We then derived Ψ,Φ angles from tau core PDBs (*Cal*) using the program PyRosetta (Chaudhury et al., 2010). Finally, we calculated the sum of the absolute per residue difference in dihedral angles, namely

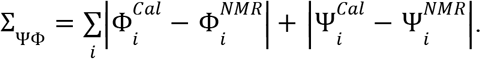

**Table S1** also shows the results of this calculation. In this case PDB entry 7p6c, a structure extracted from limbic-predominant neuronal inclusion body tauopathy (Shi et al., 2021), gave the lowest overall difference. **Fig. 4B** illustrates the Ψ,Φ angles calculated from the NMR chemical shifts (black) and 7mkg (fuchsia).

Considering that the AD fold cryo-EM structure of recombinant tau(297-391), formed under very similar conditions to ours, should be the “correct” answer, it is surprising that our three metrics result in good scores for non-AD tau core structures (see **Fig. 5A**). Although none of our metrics are perfect, Σ_A_ seems to produce the most reliable results with 8 of the 10 top scoring structures having AD folds in contrast to only 2 of the 10 bottom scoring structures. When looking at all three metrics, PDB entry 7mkg has the overall best fit with the lowest Σ_CS,_ second lowest ΔS, and 4th lowest Σ_ΨΦ_.

**Figure 5.**
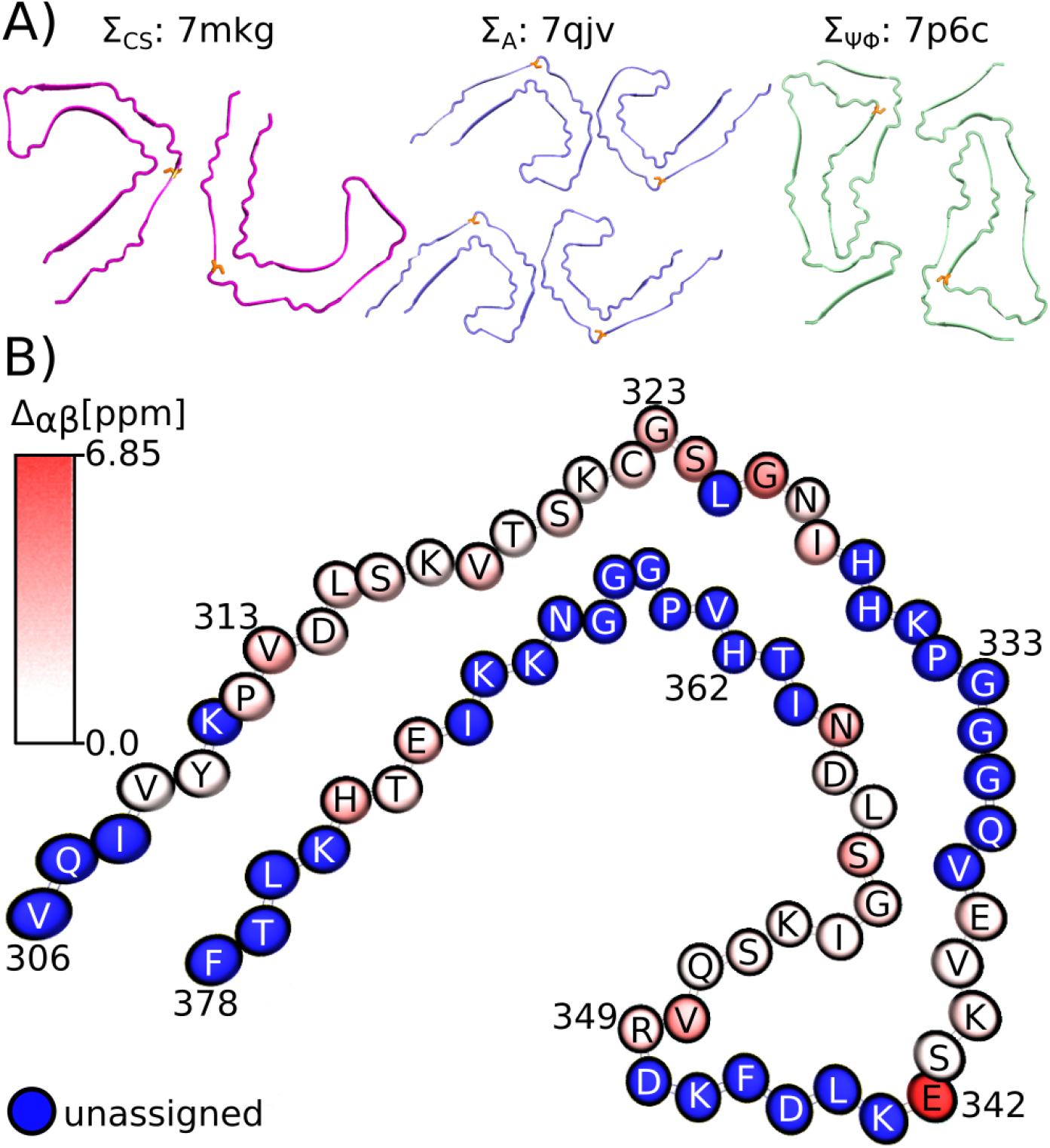
Fibril structures best fitting to NMR assignment. A) top view of fibril structures that fit best using the Σ_CS_ metric (7mkg), the Σ_A_ metric (7qjv, 7p6d not shown), and the Σ_ΨΦ_ metic (7p6c). Cys 322 is highlighted in orange. B) Birds-eye view on chain A of PDB entry 7mkg. Assigned residues are colored according to the per residue Cα and Cβ chemical shift difference calculated using the program ShiftX2 i.e. 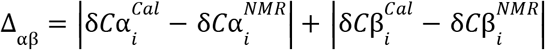 Unassigned residues are shown in blue. Residue number and type are indicated. Panel B was made with the help of the program VMD.

**Fig. 5B** illustrates the location of the residues in our assignment on the AD fold of tau (7mkg) and indicates the per residue difference between the measured (NMR) and calculated Cα and Cβ chemical shifts i.e.

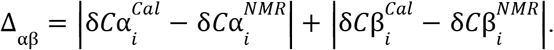

Overall, the agreement is very good as seen in **Fig. 4A**, with the biggest difference found for E342 followed by G226 and S322.

## Discussion

Cysteine residues in in-register parallel β-sheet structures result in a continuous array of thiols whose oxidation state is likely to influence both the formation and structure of the fibril. We previously showed that the oxidation state of C322 has important consequences for tau(297-391) filament forming kinetics. Electron micrographs of these two filament types had shown a similar appearance with the non-DTT filaments being much shorter (Al-Hilaly et al., 2017). Here, we followed up on these investigations using solid-state NMR on tau(297-391) filaments formed both in the presence and absence of DTT as a reducing agent. We found these structures to be remarkably different. For tau(297-391)+DTT, static well-ordered filaments were formed with few dynamic residues indicating that a large fraction of the tau(297-391) monomer is part of the fibril core. In contrast, tau(297-391) filaments formed in the absence of DTT were much less ordered as indicated by the relatively broader CP spectrum of these filaments. In addition, the relatively more intense RINEPT spectra indicated that a much larger portion of the sample was dynamic. This could either be the result of a smaller fibril core and a larger fraction of tau(297-391) remaining intrinsically disordered in the filament state, or a larger fraction of tau(297-391) staying as dimers in solution in equilibrium with the filament. These results support our previous hypothesis that disulfide-based tau dimerization is diametral to fibril formation which is in line with findings indicating that disulfide formation results in off-pathway aggregates (Walker et al., 2012). However, there are several studies arguing that disulfide formation is an important step in tau fibril formation (Schweers et al., 1995; Barghorn and Mandelkow, 2002; Furukawa et al., 2011; Kim et al., 2015) and that the inhibitory effect of methylthioninium chloride is the prevention of the formation of on-pathway disulfide bonds (Akoury et al., 2013; Crowe et al., 2013). In contrast to our present work, these somewhat contradictory results were obtained on full-length tau, which has multiple cysteine residues, aggregated in the presence of heparin.

The use of heparin to form in vitro PHF also complicates the comparison of our solid-state NMR assignment with other assignments of tau and tau fragments. There are two solid-state NMR assignments of filaments formed by the tau fragment K19 in the presence of heparin (Andronesi et al., 2008; Daebel et al., 2012). Daebel and co-workers were able to assign residues V306-S324 and observed a doubling of the resonances close to C322 caused by the different oxidation state of this cysteine. A C322A mutant removed this splitting and improved the overall quality of the spectra (Daebel et al., 2012). Dregni and co-workers presented the assignment of 0N4R and 0N3R filaments formed in the presence of heparin (Dregni et al., 2019, 2021). They identified the static core of 0N4R filaments to roughly be where in-vivo derived fibril cores were located. However, the fibril core of 0N3R extended far beyond current tau core structures spanning from R1 until the end of the C-terminus (261-441). Although all solid-state NMR assignments of filaments formed in vitro generally agree in the overall β-sheet structure of the core, they differ significantly in extent of their assignment. Even in the regions of tau that were assigned, the assignments differ in their per-residue coverage. Interestingly, none of these assignments was able to identify any of the PGGG fragments, probably because of their increased flexibility and repetitive nature. The chemical shift and dihedral angle based comparison of our NMR assignment with cryo-EM structures of mostly patient derived tau fibril cores was not able to establish a unique link between our assignment and one known structure. This is surprising because we expected the cryo-EM structure of recombinant tau(297-391), formed under very similar conditions to ours, to be the best match (Lövestam et al., 2022). However, it is important to note that our comparison is limited by the precision of the chemical shift prediction with the program Shiftx2 (with an RMSD for ^13^C shifts in the order of 0.5 ppm (Han et al., 2011, 2)), the dihedral angle prediction with the program TalosN (with a Φ,Ψ RMSD of about 12° (Shen and Bax, 2013)), and, of cause, the quality of the respective structure and of our assignment. Nonetheless, our analysis provides support for tau(297-391) filaments made in the presence of DTT adopting the AD fold. Our data suggests that comparing the position of the kinks in the β-strands by calculating the number of residues for which ΔS has a different sign might be the best metric to compare NMR assignments with existing fibrils structures. Ultimately, this analysis highlights that the difference in fibrils strains lies in large parts in changes to tertiary contacts of very similar secondary structure elements. In other words, most tau fibril strains are very similar in where their β-strand are located but different in how the resulting β-sheets fold back onto each other.

A limitation of tau(297-391) as a tau fibril model is that it represents only the most static core region of the AD fold structure. The fibril formed from full-length isoforms includes other static portions besides the core as e.g. evidenced by additional densities in cryo-EM maps (Fitzpatrick et al., 2017) and regions with static, extended structure in the 0N3R NMR assignment (Dregni et al., 2021). In addition, full-length tau filaments include the intrinsically disordered framing sequences that are known as the “fuzzy coat” in the case of tau (Wischik et al., 1988; Wegmann et al., 2013). Nevertheless, the AD fibril core represented by tau(297-391) is the binding site of powerful inhibitors of tau aggregation such as epigallocatechin gallate (EGCG) and methylthioninium, which are also able to inhibit and disaggregate preformed tau filaments (Wischik et al., 1996; Seidler et al., 2020). Identifying the binding sites of potential drugs and biomarkers provides an important step towards finding new therapeutic and diagnostic tools in AD and other tauopathies. The NMR assignment presented here could form the basis of NMR-assisted screening and characterization of such drug candidates and biomarkers.

## Data Availability Statement

Resonance assignments and time domain data will be available under BMRB ID 51483.

## Author contributions

YA optimized and prepared the samples and contributed to managing the project. CH packed the NMR sample and measured NMR data. JER expressed and purified ^13^C,^15^N labeled tau(297-391). CRH, JMDS and CMW contributed to the conception of the study and reviewed drafts of the manuscript. LCS managed the project, co-conceived the idea and contributed to writing the manuscript. ABS measured, processed, and analzyed NMR data and wrote the manuscript. All authors read and approved the final manuscript.

