## Supplemental Table and Figures for "Solid-state NMR of paired helical filaments formed by the core tau fragment tau(297-391)"

**Table S1: Comparison of NMR chemical shift assignment with existing tau fibril structures indicates that dGAE fibrils adopt the AD fold.** The following metrics were calculated for each of the structures identified by their PDB identification codes: Sum of the absolute per residue *δCα* and *δCβ* chemical shift differences between solid-state NMR assignment (*NMR*) and chemical shifts calculated using ShiftX2 [1] (*Cal*) over every assigned residue *i*

$$\Sigma_{CS} = \sum_i \left|\delta C\alpha_i^{Cal} - \delta C\alpha_i^{NMR}\right| + \left|\delta C\beta_i^{Cal} - \delta C\beta_i^{NMR}\right| .$$

Number of residues for which the difference in secondary chemical shift between Cα and Cβ (*ΔS*=*ΔδCα*-*ΔδCβ*) has a different sign when calculated using ShiftX2 as compared to the NMR assignment.

$$\Sigma_A = \sum_i A_i \text{ with } A_i := \left\{1\ if\ (\Delta S_i^{Cal}\ \cdot\ \Delta S_i^{NMR}) <\ 0,\ 0\ if\ (\Delta S_i^{Cal}\ \cdot\ \Delta S_i^{NMR}) \geq\ 0\right\}$$

Sum of the absolute per residue Ψ, Φ differences as calculated from the solid-state NMR assignment using TALOS-N (*NMR*) and calculated from the PDB structure (*Cal*)

$$\Sigma_{\Psi\Phi} = \sum_i \left|\Phi_i^{Cal} - \Phi_i^{NMR}\right| + \left|\Psi_i^{Cal} - \Psi_i^{NMR}\right|.$$

Best (i.e. minimal) values for each metric are underlined and the table is sorted according to $\Sigma_A$. In addition, the table lists the fibril source and fold type where the abbreviations are defined as follows: AGD, argyrophilic grain disease; PrP-CAA, PrP cerebral amyloid angiopathy; AD, Alzheimer's disease; PART, primary age-related tauopathy; LNT, limbic-predominant neuronal tauopathy; CTE, chronic traumatic encephalopathy; GSS, gerstmann-sträussler-scheinker disease; GGT, globular glial tauopathy; CBD, corticobasal degeneration; PSP, supranuclear palsy.

| PDB | $\Sigma_{CS}$ [ppm] | $\Sigma_A$ | $\Sigma_{\Psi\Phi}$ [deg] | Disease / Source | Fold |
|---|---|---|---|---|---|
| 7p6d | 65.8414 | <u>4</u> | 3648 | AGD | AGD |
| 7qjv | 73.6912 | <u>4</u> | 3361 | Recombinant | AD / Quadruple |
| 7mkg | <u>62.9429</u> | 5 | 2848 | PrP-CAA | AD / Straight |
| 5o3o | 74.1779 | 5 | 3146 | AD | AD / Paired |
| 7mkf | 71.8944 | 6 | 2883 | PrP-CAA | AD / Paired |
| 7nrs | 74.7738 | 6 | 2986 | PART | AD / Straight |
| 7nrt | 74.9086 | 6 | 2987 | PART | AD / Straight |
| 7nrx | 83.0367 | 6 | 2972 | AD | AD / Straight |
| 5o3t | 88.8657 | 6 | 3827 | AD | AD / Straight |

| | | | | | |
|---|---|---|---|---|---|
| 7p6e | 97.3397 | 6 | 3786 | AGD | AGD |
| 7p6c | 69.0672 | 7 | 2661 | LNT | LNT |
| 7p6b | 73.1256 | 7 | 2723 | LNT | LNT |
| 6nwp | 75.0648 | 7 | 3314 | CTE | CTE |
| 7nrq | 76.0444 | 7 | 3093 | PART | AD / Paired |
| 7mkh | 76.1057 | 7 | 2916 | GSS | AD / Paired |
| 7p67 | 83.657 | 7 | 3466 | GGT | GGT |
| 6tjo | 94.4223 | 7 | 3707 | CBD | CBD |
| 7nrv | 73.4251 | 8 | 2972 | AD | AD / Paired |
| 7p68 | 77.7252 | 8 | 3010 | GGT | GGT |
| 5o3l | 95.7989 | 8 | 3454 | AD | AD /Paired |
| 6tjx | 96.9895 | 8 | 3474 | CBD | CBD |
| 7p6a | 76.5164 | 9 | 2754 | LNT | LNT |
| 7p65 | 82.3178 | 9 | 3018 | PSP | PSP |
| 7p66 | 89.1693 | 11 | 2963 | GGT | GGT |
| 6nwq | 90.8825 | 11 | 3314 | CTE | CTE |

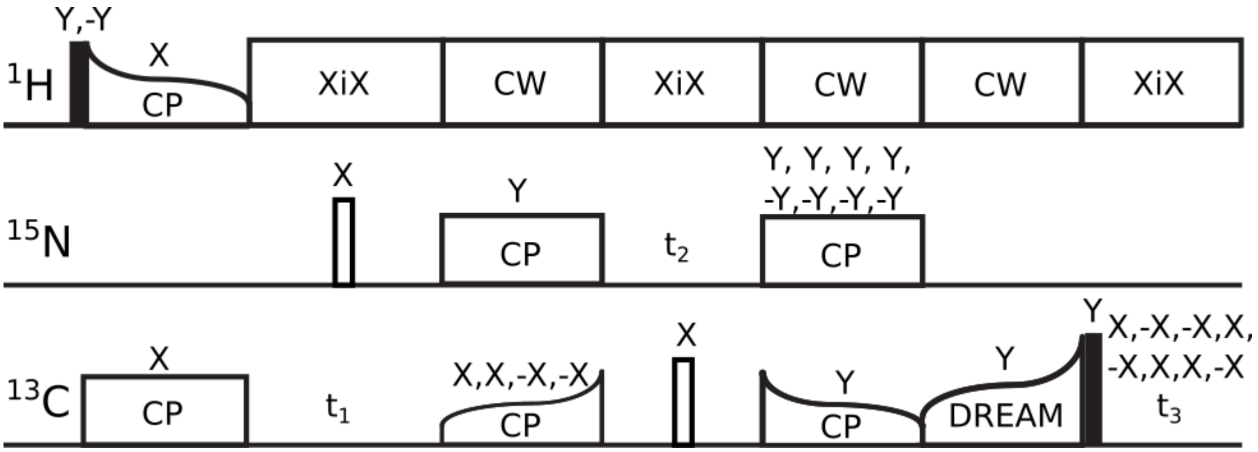


**Figure S1:** Pulse sequence for 3D NCAcoCA experiment. 90° and 180° pulses are shown as black and white rectangles, other pulse sequence elements are annotated and phases of individual pulses are given except for decoupling elements.

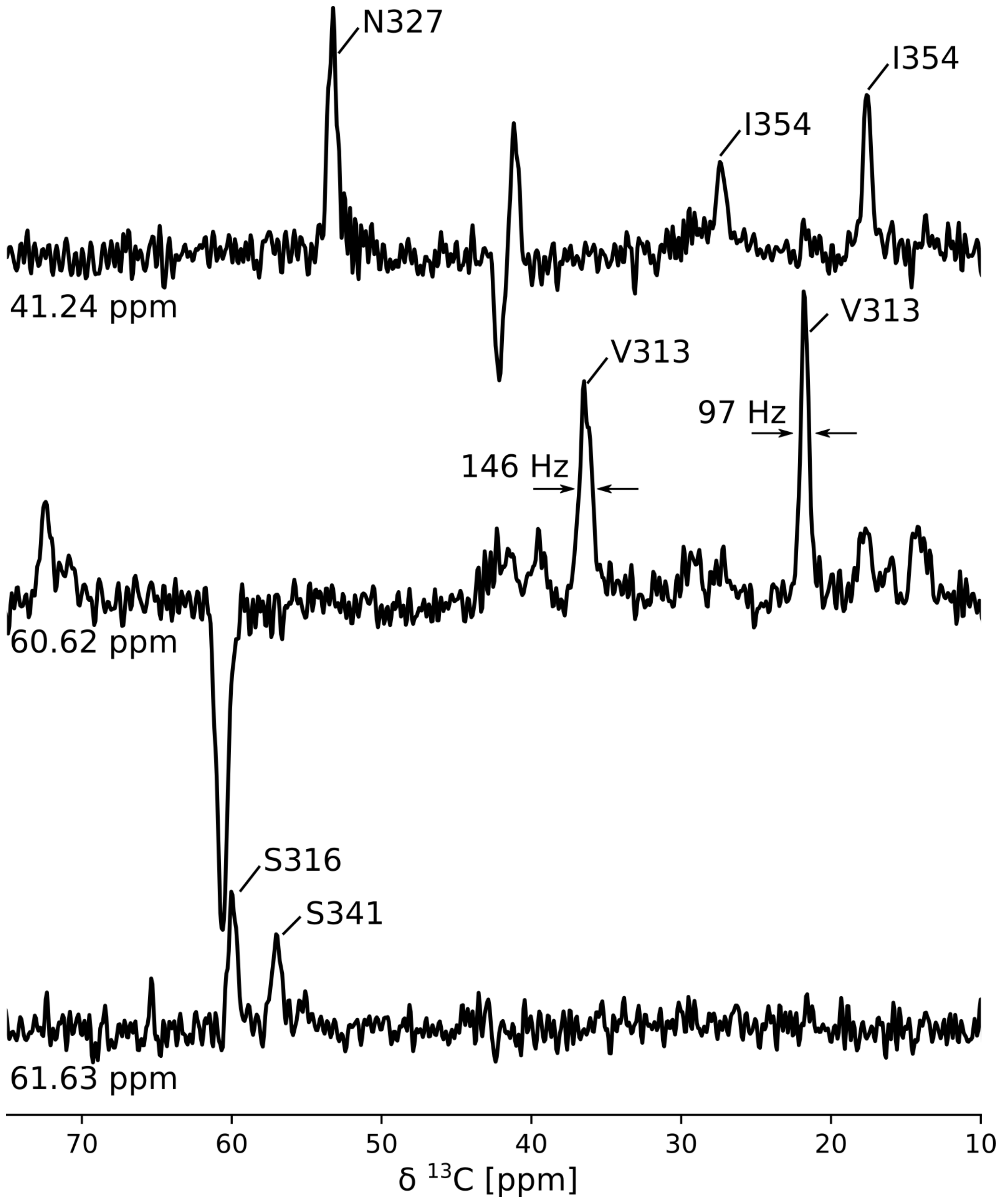


**Figure S2: 1D slices through 2D DREAM spectrum illustrate quality of dGAE+DTT sample.** 1D slices through the 2D DREAM spectrum shown in Fig. 2A, which was processed without window function in the direct dimension for the present figure. PPM positions in the indirect frequency domain, assignments of cross peaks, and linewidths for V313 Cβ and Cγ resonances are indicated.
